# Intra- and Interspecific Divergence of FXPRLamide Neuropeptide-Receptor Interactions in flies (Diptera)

**DOI:** 10.1101/2025.07.25.666740

**Authors:** Sarah M. Farris

## Abstract

Pyrokinin (PK) neuropeptides are characterized by a conserved C-terminal FXPRLamide motif and modulate a range of physiological functions and behaviors in species spanning the Eumetazoa. The *pban* gene is conserved across the insects and encodes pheromone biosynthesis activating neuropeptide (PBAN) and 1-3 additional pyrokinins. The *pban* gene of the basal Diptera resembles that of other insect Orders while in more derived Diptera (where it is referred to as the *hugin* gene) the PBAN peptide coding sequence is absent and another peptide, hugin, serves novel functions in the fruit fly *Drosophila melanogaster*. In this paper, structural models are generated for the pyrokinins PBAN and γ-SGNP and their receptor PBANR of the basal species *Aedes aegypti*, and hugin and its receptor PK2_1 of the more derived *Drosophila melanogaster.* The binding pockets for all three peptides overlap even when amino acids are non-synonymous between the two species. Peptide C-terminal core sequences preferentially bind more conserved transmembrane regions of the receptor while the variable N-terminal peptide residues are relegated to less conserved sites in the extracellular loops. The PBAN peptide forms rigid secondary and tertiary structures with its long N-terminus, binding unique amino acids in the extracellular loops of PBANR, providing a basis for functional differentiation from the short and flexible γ-SGNP peptide. A kink introduced into the full peptide backbones by proline residues directs the N-terminus along the surface of the ECLs, suggesting roles for secondary and tertiary structure in peptide recognition. *D. melanogaster* hugin binds residues that contact PBAN in *A. aegypti,* suggesting that loss of PBAN in higher Diptera facilitated divergence of peptide-receptor interactions. This data provides a potential basis for differing roles of multiple peptides in *A. aegypti*, and coevolution of ligand and receptor in *D. melanogaster* that may have facilitated functional evolution.

## INTRODUCTION

Neuropeptides are evolutionarily ancient signaling molecules that function as juxtacrine, paracrine, or endocrine modulators of physiology and behavior. Many neuropeptide families and their G-protein coupled receptors (GPCRs) are conserved across the Metazoa, underscoring the importance of neuropeptide signaling [1]. However, diversification of neuropeptides and their cognate receptors reflects their adaptability to physiological and behavioral requirements of diverse ecological niches encountered over millions of years of animal evolution.

### The PRXamide family of neuropeptides

The PRXamides amply illustrate the effects of selection on neuropeptide evolution. Members of this family have been identified in the genomes of animals across the Metazoa based on their shared C-terminal sequence, in which the terminal “X” is typically an uncharged residue [2]. PRXamides are broadly distributed across the protostomes and are considered homologous to deuterostome Neuromedin U, with more distant relationships to TRH and ghrelin [3,4]. Within the non-deuterostome Metazoa, PRXamides and their receptors have been recovered from species representing he Euarthropoda, Mollusca, Annelida, Nematoda and the Cnidaria [1,5–8].

In the Euarthropoda, genes encoding three classes of PRXamides have been identified: *ecdysis-triggering hormone* (*eth*), *capability* (*capa*), and pheromone biosynthesis-activating neuropeptide (*pban*, or *hugin* in *Drosophila melanogaster*); for review see [2]. ETH peptides regulate ecdysis and are conserved across the Panarthropoda [9]. Capa neuropeptides including perviscerokinins (PVKs) and the diapause hormone DH-1; PVKs have myotropic and osmoregulatory functions while DH-1 regulates diapause onset [10–14]. The *pban* gene has been isolated from species of Chelicerata and Pancrustacea [15–18]. In insects the gene encodes a variable number of peptides with a C-terminal FXPRL core motif including an additional diapause hormone (DH-2 or PK1) and 1-4 pyrokinins (PK2s) including PBAN. The PK2s are inconsistently named in the literature but this account will use the terms SGNP-α, -β and -γ (subesophageal ganglion neuropeptide) following the nomenclature established in the Lepidoptera. α-SGNP appears mostly restricted to the Lepidoptera while β-SGNP, PBAN, and γ -SGNP peptides are present in many species (for reviews see [19–22]).

### FXPRLamide receptors

All pyrokinins bind Class A G protein coupled receptors (GPCRs), with separate receptors for DH (DHR) and for PBAN and the SGNPs (PBANR or PK2R). While each peptide typically binds its cognate receptor with high affinity, cross-reactivity is common at physiological concentrations [23–31]. Additional diversity of cellular responses is provided by tissue-specific expression of multiple receptor splice isoforms [27,31–33] or duplication of the receptor as observed in the higher Diptera [34,35].

### Roles of neuropeptides encoded by the PK/PBAN gene

As the name suggests, PBAN is a regulator of sex pheromone biosynthesis and/or release [36–39] although functional divergence is apparent in other species. Supporting a function as a general regulator of intraspecific communication, *pban* gene encoded neuropeptides initiate aggregation pheromone synthesis in the western flower thrips *Frankliniella occidentalis* [40] and trail pheromone synthesis in the fire ant *Solenopsis invicta* [41]. The trail pheromone gland is derived from Dufour’s gland [42], which produces several intraspecific communication signals including sex pheromone in Hymenoptera [43,44]. Intriguingly, transformation of an entomopathogenic fungus with the *Solenopsis invicta* coding sequence for β -SGNP peptide increased virulence of a fungal insecticide and associated behavioral abnormalities in the ant [45]. Additional roles for peptides encoded by the *pban* gene, including DH2, include regulating cuticle pigmentation [28,46] and timing of life history transitions such as pupariation and diapause [47–50]. In the fruit fly *Drosophila melanogaster* the *pban* gene homolog *hugin* encodes two peptides, hugin and γ-hugin. Hugin is likely homologous to γ-SGNP and plays complex roles in the regulation of feeding associated locomotor behavior under circadian regulation [51–55].

The functions of the remaining peptides encoded by the *pban* and *hugin* genes are poorly understood. In *D. melanogaster* γ-hugin (likely homologous with β-SGNP) possesses a canonical FXPRLamide motif and activates PK2_1 receptors *in vitro* even though the peptide is undetectable *in vivo* [34,56]. In other species SGNP peptides bind PBANR as efficiently as PBAN itself and may bind DHR at lower affinities; likewise, DH2 can bind PBANR in some instances [23,29,30,48,57–60]. The benefits conferred by such receptor-ligand promiscuity are unknown.

### Conservation and divergence of PK/PBAN neuropeptides in Diptera

The *pban* gene in the basal Diptera encodes a prepropeptide with an ancestral complement of neuropeptides: DH2, PBAN and β- and γ-SGNPs. DH2 is lost at the origin of the Neodiptera and PBAN is absent in the Syrphidae + Schizophora. A conserved γ-SGNP sequence SVXPFKPRL (aka hugin) is characteristic of the Brachycera as is a more variable β-SGNP, aka γ-hugin [22,61]. It is well established that conserved neuropeptide families coevolve with their receptors [1], so it is likely that the loss and divergence of FXPRL neuropeptides such as PBAN in the Diptera are associated with corresponding changes in their cognate receptors. In this study, multiple approaches are used to investigate peptide-receptor interactions in the yellow fever mosquito *Aedes aegypti* and the fruit fly *Drosophila melanogaster* to identify such coevolutionary signatures. Sequence alignments, binding pocket predictions and docking models reveal conserved and unique peptide binding sites on their cognate receptors and distinct secondary and tertiary structures, all providing a potential basis for modulation of receptor activation and divergence of function while maintaining recognition of the FXPRL core motif.

## METHODS

### Gene sequences and binding pockets

The translated *Aedes aegypti* PBAN (GenBank accession #XM_001662162 and PBANR (GenBank accession #KC155994)[62] and *Drosophila melanogaster* hugin (CG6371, accession #AJ133105) [61] and PK2_1R (CG8784, accession #AF522189)[35] were aligned using Clustal Omega [63]. The *D. melanogaster* genome also contains a second PK2 receptor (PK2_2, CG8795) with 58.8% identity to CG8784. Both receptors are strongly activated by hugin [35] although also see [34], but only PK2_1 has been shown to mediate physiological functions of hugin in *D. melanogaster* [64] and was thus chosen for investigation in this study. Binding pockets of each receptor were identified from sequence data using PrankWeb [65], and mapped onto sequence alignments. Predicted amino acid contacts between ligand and receptor using the methods described in the following section were also mapped onto sequence alignments and onto receptor secondary structural models generated using the Protter tool [66].

### Modeling peptide-receptor binding

Receptor structure models were retrieved from the AlphaFold protein structure database (*A. aegypti* PBANR AF-V9P4J1-F1-v4; *D. melanogaster* PK2_1R AF-Q9VFW5-F1-v4)[67]. Most of the flexible, disordered extracellular N-terminus of each protein was removed from these models to facilitate peptide docking Kawai et al (2014)[68]. Structure models of each peptide were generated from the active peptide sequences (*A. aegypti* PBAN DASSSNENNSRPPFAPRL; γ-SGNP NLPFSPRL) and *D. melanogaster* hugin (SVPFKPRL) using PepFold4 [69]. Only those peptide models that formed a β-turn between the X and L residues of the core sequence, required for receptor activation, were selected for docking analysis [70–72]. Two tools were employed to assess interactions between individual amino acids of peptides and receptors. MdockPep generated docking predictions using receptor structural models and peptide sequences [73] while predictions made from structural models of both peptides and receptors were generated using ClusPro 2.0 [74]. Predicted ligand-binding domains were visualized using Clustal Omega and Protter as described above. Only those models in which the core sequence penetrates the binding pocket were selected for further investigation [75,76]. Docked peptides and receptors were imported into UCSF Chimera X [77] for visualization, identification of amino acid contacts, presence and location of β-turn hydrogen bonds and matchmaking between isolated and full peptide core structures.

## RESULTS

### Comparisons of the *A. aegypti* PBANR and *D. melanogaster* PK2_1R proteins and binding pockets

Sequence alignment of *A. aegypti* PBANR and *D. melanogaster* PK2_1R proteins revealed substantial homology with an overall identity of 43.49% (Fig 1, S1 Table). Lowest identity occurs in the extracellular N-terminus (18.3%), the intracellular C-terminus (23.4%), and the third intracellular loop (ICL3; 34.5%). The second intracellular loop (ICL2; 95%) and transmembrane domains 4–7 (71.43–79.17%) are most highly conserved, consistent with their known roles in G-protein coupling and signal transduction [78,79]. Binding specificity of peptide ligands to their cognate GPCRs occurs mostly via interactions with the three extracellular loops (ECLs) and extracellular-facing residues of the transmembrane domains [68]; these regions have low to intermediate similarity between the two species. Binding pocket predictions made using PrankWeb corroborate these features of PBANR and PK2_1R.

**Fig 1.**
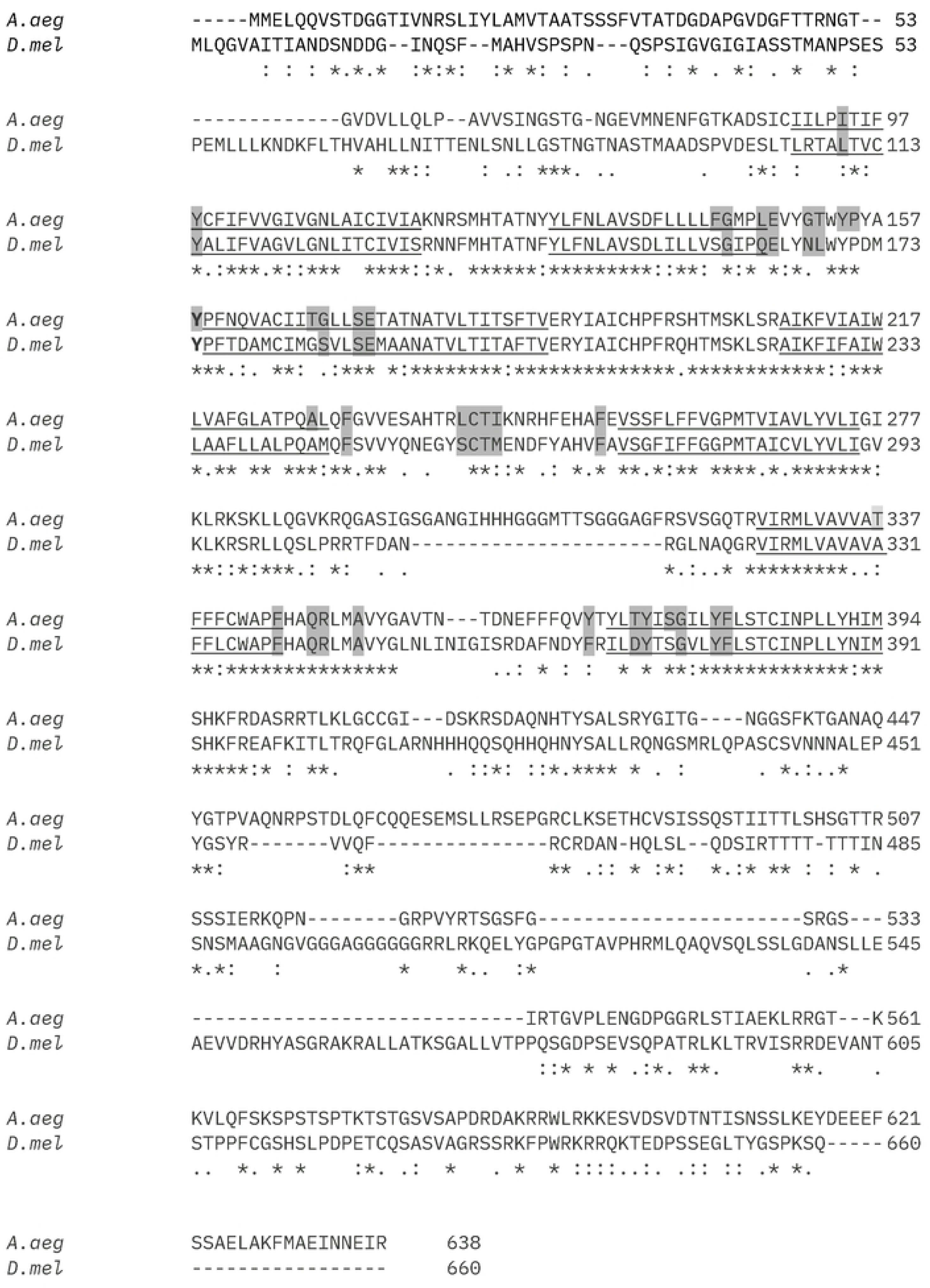
Sequence alignment of *A. aegypti* PBANR and *Drosophila melanogaster* PK2_1R. Transmembrane regions are underlined and binding pocket residues predicted by PrankWeb are indicated in dark grey. Annotations below alignment: * conserved, : conservative mutation,. semi-conservative mutation, no symbol- non-conservative mutation.

The structure of the *A. aegypti* PBAN, γ-SGNP and *D. melanogaster* hugin prepropeptides are shown in Fig 2A. PBAN is characterized by a FAPRL core motif and long N-terminal sequence (DASSSNENNSRPP); γ-SGNP consists of a FSPRL core and short N-terminus (NLP). Hugin, like γ-SGNP, has a three amino acid N-terminus (SVP) but a FKPRL core motif. MDockPep predictions of interactions between peptide core motif and N-terminal residues with specific receptor loci (Fig 2B) are consistent with binding pocket predictions in Fig 1. Receptor binding by FXPRL core sequences (shaded) is frequently localized to the transmembrane regions (underlined) while binding sites of the peptide N-terminus (bold) often occur in the ECLs and receptor N-terminus, although there is substantial overlap between the two (shaded + bold). Overall ligand-receptor interactions between the Diptera peptides and receptors, *Bombyx mori* PBAN and PBANR and with human NMU/NMS to NMUR1 and NMUR2 (the vertebrate homolog of insect pyrokinins/PKRs) reveal conserved binding sites especially in the ECLs, TM3 and TM7 (S2 Fig). [75,76,80]

**Fig 2.**
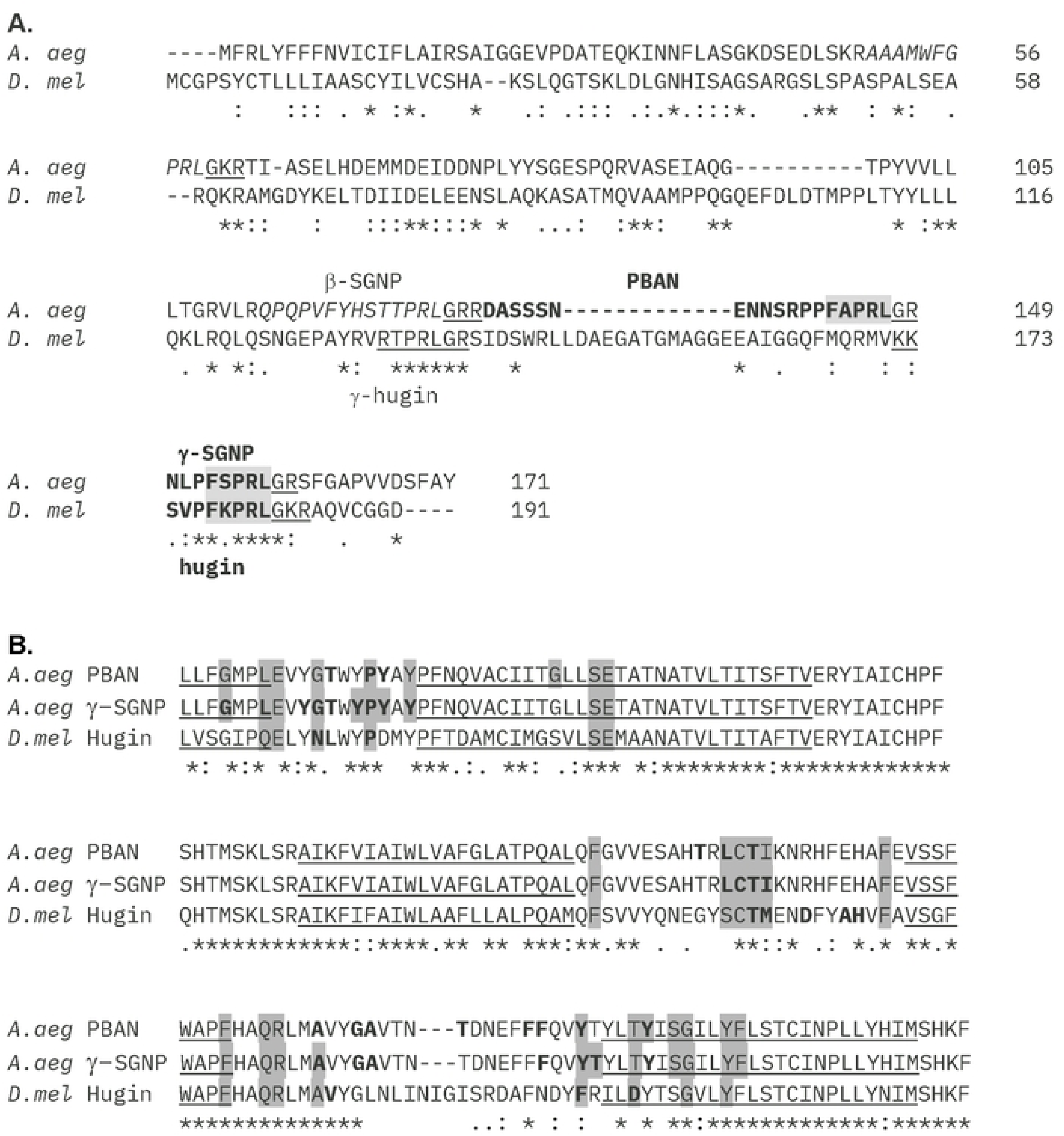
A. Aligned sequences of the *A. aegypti* PBAN and *D. melanogaster* hugin prepropeptides. A. PBAN, γ-SGNP (Q16N80) and hugin (NP_524329) peptide sequences are bolded and labeled. Core motifs are indicated by a shaded box while amidation and peptidase cleavage sites are underlined. Locations of additional pyrokinin peptides are highlighted in grey. B. Peptide binding residues of *A. aegypti* PBANR (accession number AGT80483) and *D. melanogaster* PK2_1R (NP_731790) as predicted by MDockPep. Grey boxes indicate core sequence binding (FXPRL), bold indicates binding by N-terminal residues of the peptide, both indicate binding by core and N-terminal residues. Only extracellular loops and nearby transmembrane regions (underlined) are shown. Annotations below alignment: * conserved, : conservative mutation,. semi-conservative mutation, no symbol- non-conservative mutation.

ClusPro docking predictions using structural models of the peptides and their receptors revealed substantial overlap of ligand-receptor binding patterns overlaid by peptide- and species-dependent variation. The snake plot in Fig 3A summarizes total ligand-receptor interactions of the three peptides studied here, using PBANR of *A. aegypti* as a template. A total of 35 receptor amino acids interact with all three ligands (light blue), with some engaging with the same amino acids in all ligands of both species (S3 Fig) even if a non-synonymous amino acid substitution occurs between *A. aegypti* and *D. melanogaster* (red outline). This exceeds the number of receptor contacts predicted by MDockPep, reflecting the multiple and somewhat variable valid structural models and predictions generated by PepFold4 and ClusPro. *A. aegypti* PBAN has the most unique receptor binding interactions (13 residues; Fig 3A grey), primarily involving non-synonymous sites in the ECLs. Hugin makes six unique interactions (yellow), four of which are with non-synonymous residues, and γ-SGNP makes three unique interactions with conserved receptor residues (magenta). Hugin shares six binding sites with PBAN (orange; four non-synonymous) and three with γ-SGNP (green; all conserved).

**Fig 3.**
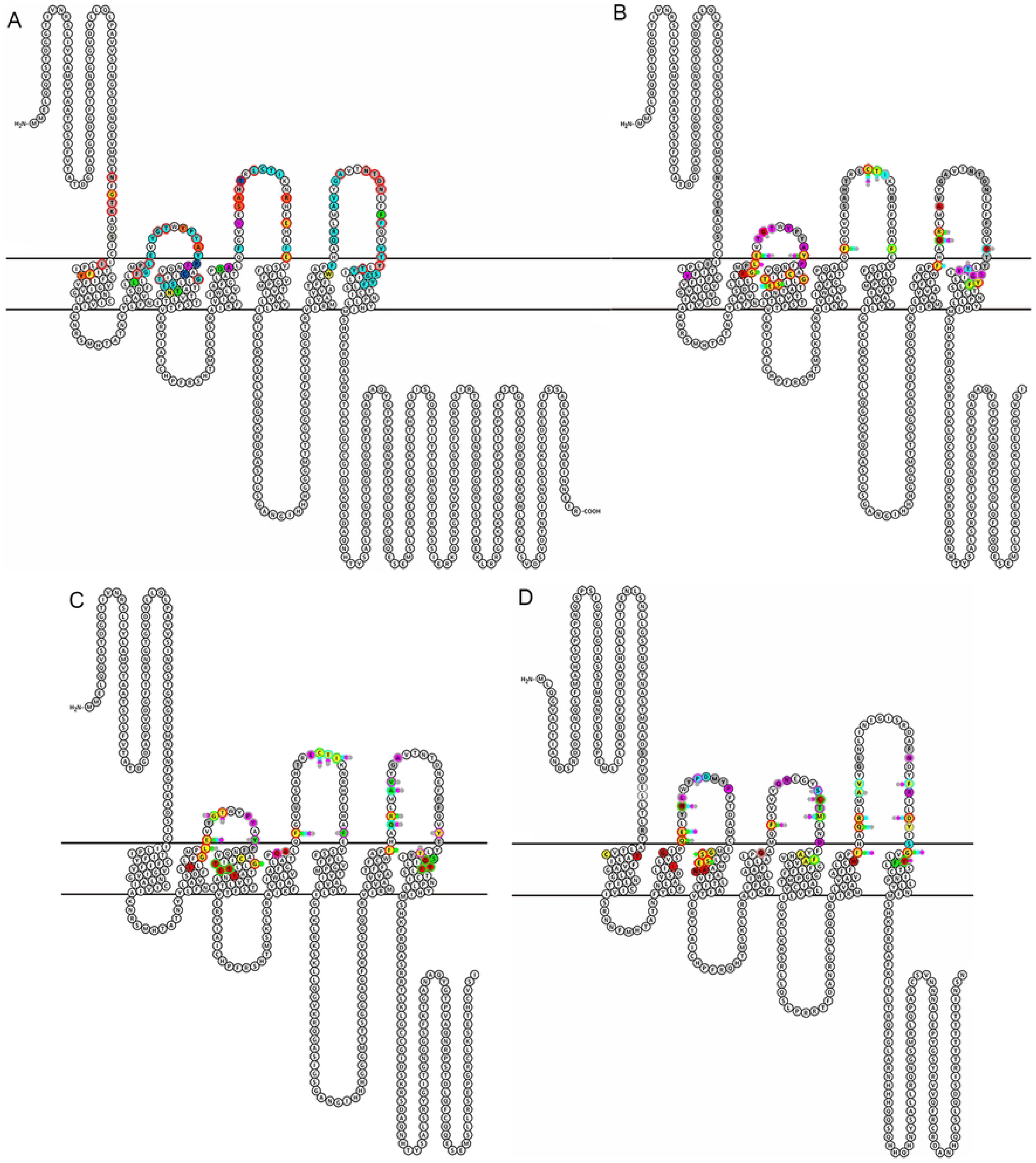
Snake plots of ligand-receptor docking predictions. A. Summary of *A. aegypti* PBAN, γ-SGNP, and *D. melanogaster* hugin peptide binding sites using the *A. aegypti* PBANR as a template. Light blue, receptor residue bound by all peptides; grey, bound only by *A. aegypti* PBAN; magenta, bound only by *A. aegypti* γ-SGNP; dark blue, bound by both *A. aegypti* PBAN and γ-SGNP; yellow, bound only by *D. melanogaster* hugin; green, bound by both *A. aegypti* γ-SGNP and *D. melanogaster* hugin; orange-bound by both *A. aegypti* PBAN and *D. melanogaster* hugin. Red circles-weakly conservative or non-conservative mutation between *A. aegypti* and *D. melanogaster*. B-D. Binding of peptide ligands to their cognate receptor by individual peptide residue. B. *A. aegypti* PBAN/PBANR. **C.** *A. aegypti* γ-SGNP/PBANR. D. *D. melanogaster* hugin/PK2_1R. Grey, N-terminal peptide amino acids: FXPRL residues magenta, Phe (F); blue, X; green, Pro(P); red, Arg (R); yellow, Leu (L).

Analysis of specific ligand-receptor interactions reveals consistent features across peptides and species (Fig 3B-3D; S4 Table). Concordant with the MDockPep data, individual neuropeptide residues bind multiple receptor residues, and vice versa. As previously described, FXPRL core motifs, especially the Pro-Arg-Leu residues (green, red and yellow respectively) of all peptides interact frequently with transmembrane domains. Phe and X residues (magenta and blue) of the core motif often share binding sites in the ECLs with the N-terminal domains of the peptides. Interactions between peptide N-termini (grey) and distal ECLs more frequently involve a single peptide residue with the exception of a Cys-containing β-turn motif in ECL2 characteristic of GPCRs that binds nearly all core motif and N-terminal peptide residues [81]. In all peptides the variable “X” of the FXPRL core sequence forms the fewest bonds with the receptor and is almost entirely restricted to the ECLs (Fig 3B-3D; S4 Table). The core motif Pro residue of PBAN and hugin binds infrequently but in γ-SGNP makes additional contacts in TM3 and ECL3. The core motif C-terminal Leu of hugin interacts more frequently with ECL3 than is observed for the other peptides. The PBAN N-terminus binds PBANR almost exclusively at the ECLs and has the most binding sites, in keeping with the long (13 amino acid) N-terminus compared with those of the other two peptides (3 amino acids each).

### Ligand-receptor structural models and docking predictions

Docking predictions generated by ClusPro for PBAN, γ-SGNP and hugin revealed differences in secondary and tertiary peptide structures and their interaction with their cognate receptors (Fig 4, S5 Fig). Fig 4A-B, E-F, and I-J depict the top two ClusPro predictions using the top two PepFold4 models of PBAN, γ-SGNP and hugin respectively. The figures illustrate the greater length of the PBAN peptide allowing interaction with distal portions of the ECLs in comparison with the compact γ-SGNP and hugin peptides. Ribbon models in Fig 4C-D, G-H and K-L combine all valid peptide models and their docking predictions. There is some variability among the models, especially γ-SGNP and hugin but also in the orientation of the core motif β-turn in the binding pocket of all three peptides. PBAN is unusual in that its extended N-terminus forms a rigid helix that rests alongside ECL3 and approaches ECL2 at its most distal (Fig 4C-D). The FAPRL core also adopts a helical structure, consistent with the helix-promoting and stabilizing properties of Ala and Pro [82–84]. PepFold4 models of γ-SGNP reveal less ordered structures, and docking predictions suggest that peptide orientation within the binding pocket is more flexible that that observed for PBAN (Fig 4E–H). The short N-terminus remains within the binding pocket but is also predicted to be capable of a wider range of receptor interactions than for PBAN. Hugin models and predictions display similar features to γ-SGNP (Fig 4I–L).

**Fig 4.**
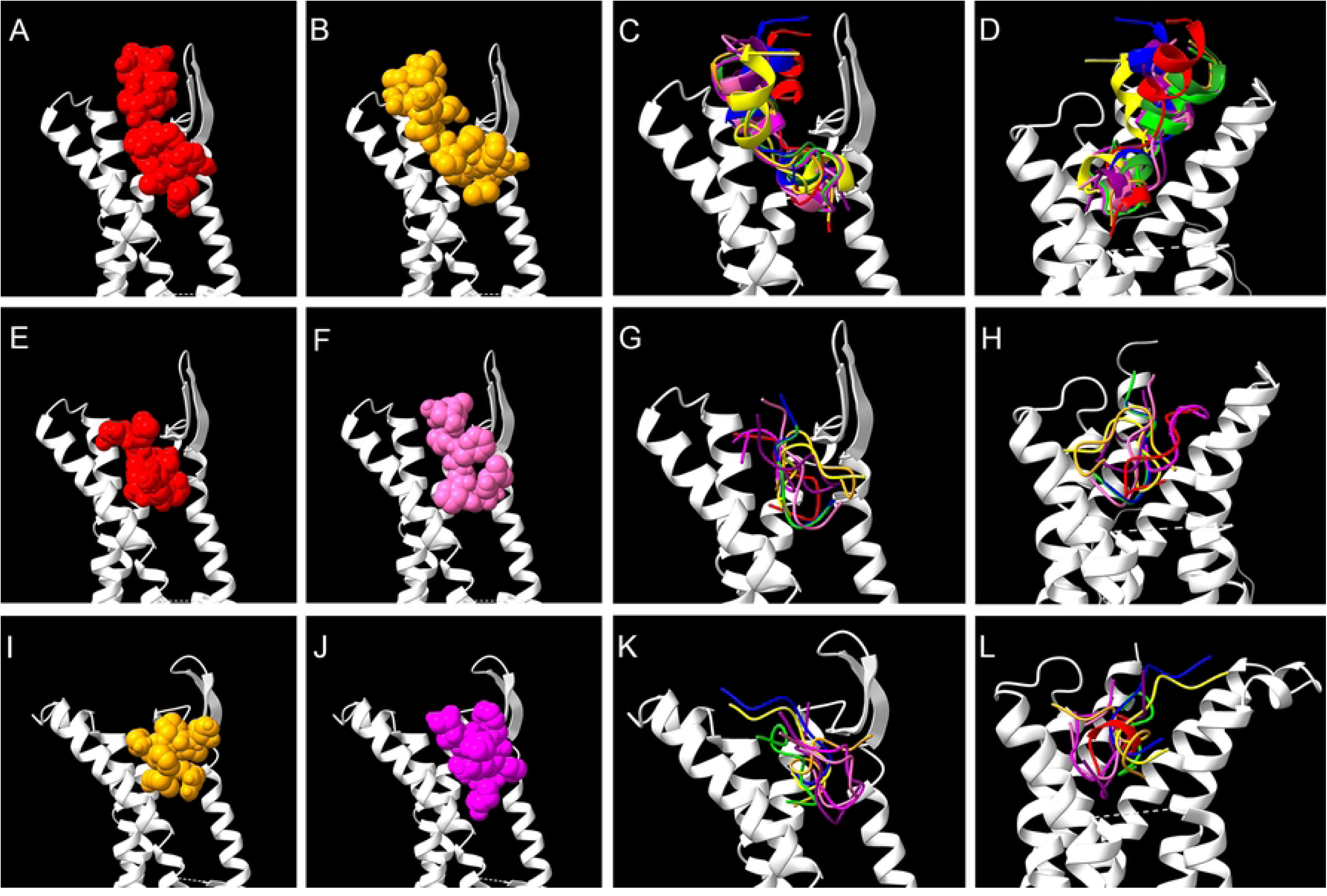
Docking predictions using peptide and receptor structural models. A-B: Top two docking predictions for *A. aegypti* PBAN to PBANR. ECL1 and N-terminus are removed to provide a clearer view of the peptide. C: Ribbon models of all valid PBAN peptide structural models and docking predictions with the N-terminus and ECL1 removed. D: Same as previous except with ECL2 removed. E-F: Top two docking predictions for *A. aegypti* γ-SGNP to PBANR. ECL1 and N-terminus are removed to provide a clearer view of the peptide. G: all valid γ-SGNP peptide docking predictions with the N-terminus and ECL1 removed. H: Same as previous except with ECL2 removed. I-J: Top two docking predictions for *D. melanogaster* hugin to PK2_1 to PBANR. ECL1 and N-terminus are removed to provide a clearer view of the peptide. K: all valid hugin peptide docking predictions with the N terminus and ECL1 removed. L: Same as previous except with ECL2 removed.

The PBAN core motif alone is sufficient to activate PBANR *in vitro*, and the isolated FAPRL core sequence interacts with the same receptor residues (yellow, Fig 5A, 5B) as that in the full peptide (blue). Arrows indicate two proline residues between the core motif and helical N-terminus in the full peptide that introduce a “proline kink” in the peptide backbone that directs the N-terminus through the center of the receptor. Structural models of bound isolated core sequences (yellow) are not aligned with the full peptide core sequence (magenta, Fig 5C). Forced alignment of the full peptide and isolated core sequences using the Matchmaker function in Chimera X dramatically reorients the full PBAN N-terminus so that instead of projecting through the center of the binding pocket it protrudes orthogonally through ECL3, directed sideways out of the binding pocket by the proline kink (green Fig 5D, 5E; arrows-proline residues). Forced alignment of the isolated (yellow) and full peptide core sequence (magenta) moves the Phe residue of the latter to nearly the same position as in the isolated core (Fig 5F, arrows). The proline kink in this model directs the (Fig 5E) median backbone angle of the Phe-Pro-Pro sequence is 122.6° while that of Pro-Pro-Arg is 121.3° (data not shown). Docking predictions for the isolated core sequence cannot incorporate this bend, resulting in binding interactions with the pocket that would not be permissible for the full peptide. Fig 5G depicts a consensus alignment of peptide of full PBAN and isolated core motif binding to PBANR, color coded by peptide residue. While the overall pattern of interactions is similar, individual amino acids of the full peptide core sequences make more frequent contacts that often bind more than one peptide residue. The distribution of binding sites across the receptor also differs. For example, Phe of the FAPRL core motif is adjacent to ECL1 in the full peptide but more centrally positioned in the isolated core sequence, resulting in fewer interactions between Phe and ECL1 in the latter. Similar analyses were conducted for γ-SGNP and hugin (S6 Fig). As with PBAN, these peptides have a Pro residue immediately before the core sequence, with the resulting kink directing the N-terminus along the inner surface of ECL3 but orthogonally through the ECL when isolated and full peptide core sequences are forced to align. The median backbone angle around the proline residue of hugin is close to that of PBAN (V-P-F; 124.0°), that of γ-SGNP is smaller (L-P-F; 115.7°; data not shown).

**Fig 5.**
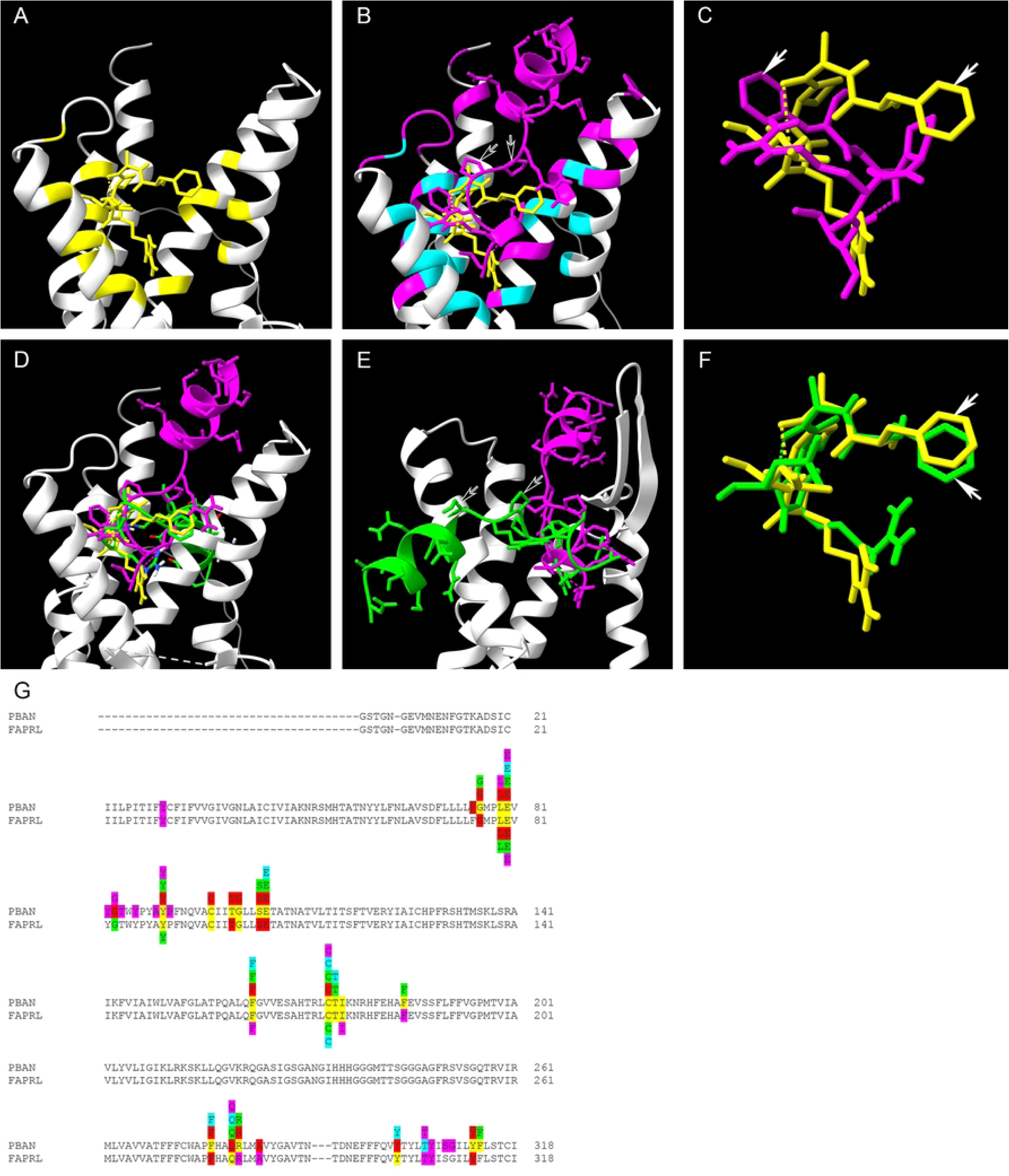
Docking of PBAN core motif and the full *A. aegypti* PBAN peptide to PBANR before and after alignment of core sequences. A. Receptor residues contacted by the isolated FAPRL core sequence (yellow), with ECL2 removed for clarity. B. Isolated core motif (yellow) and full PBAN peptide (magenta) docked to PBANR. Overlap of receptor contacts between the isolated core motif and the full peptide are indicated in blue, PBAN peptide N-terminal contacts only in purple. Arrows-proline residues producing a “proline kink” in the full peptide. C. Overlap of isolated (yellow) and full peptide (magenta) core motif docking predictions (arrows). Arrows-phenylalanine residues. D. Docking of the isolated core motif (yellow) and full PBAN peptide before (magenta) and after (green) after forced alignment of the isolated core motif with that of the peptide. The helical N-terminal of the PBAN peptide is oriented orthogonally with respect to the ECLs, through the back of the page. E. Same as panel D but with the receptor rotated 180 degrees and ECL1 removed for a better view of the altered position of the peptide N-terminus. The kink produced by the double prolines (arrows) introduce a kink in the backbone that directs the helical N-terminus orthogonal to the transmembrane regions and through ECL3. F. Correspondence of the isolated core motif (yellow) and that of the full peptide (green) after forced alignment. Arrow-phenylalanine residues. G. Consensus of docking predictions for the isolated FAPRL and full PBAN core motifs binding PBANR, with colors indicating residues of each peptide. magenta, Phe (F); blue, A; green, Pro(P); red, Arg (R); yellow, Leu (L)grey-N terminal (full PBAN only). Only receptor ECL and TM binding regions are shown.

## DISCUSSION

Peptides encoded by the insect *pban* gene are conserved neuropeptides characterized by a C-terminal FXPRLamide motif. They act through G-protein coupled receptors (GPCRs) to regulate diverse physiological processes in both developing and adult insects. In basal Diptera represented here by *Aedes aegypti* (yellow fever mosquito), the *pban* gene encodes four peptides: DH2 (PK1), β-SGNP, PBAN, and γ-SGNP [22,62]. In contrast, in more derived Diptera represented here by the fruit fly *Drosophila melanogaster*, the homologous gene (*hugin*) encodes only β-SGNP (γ-hugin) and γ-SGNP (hugin) [22,61,85].

This study compares intraspecific interactions between PBAN and γ-SGNP with their receptor PBANR in *A. aegypti*, and interspecific differences between these and hugin (γ-SGNP homolog) and its receptor PK2_1 in *D. melanogaster*. Within species, PBANR binds promiscuously to PBAN gene-encoded peptides with varying affinities [24,26,30,31,34]. The data presented here provides insight into sequence and structural features of each peptide that confer unique binding interactions with PBANR that might enable peptide-specific functions. Coevolutionary changes between *D. melanogaster* hugin and PK2_1, including non-synonymous changes in receptor amino acids at binding sites shared across all peptides, acquisition of binding sites occupied by PBAN in *A. aegypti*, and fixation of the SVPFKPRL hugin (γ-SGNP) sequence are observed. The novel roles in circadian regulation of feeding and locomotor behaviors that have been uncovered in *D. melanogaster* may have evolved concurrently with such coevolutionary changes.

Sequence identity between PBANR and PK2_1R is highest within the transmembrane domains (TMs) and ICLs 1 and 2, consistent with their central role in GPCR stabilization, activation and G-protein coupling [68,78,79,86]. In contrast, extracellular loops (ECLs) which mediate ligand recognition and binding [24,81,87] show lower sequence identity, promoting ligand specificity. Nevertheless, a binding pocket comprising 35 residues in both TMs and ECLs engages all three peptides, even though some of the amino acids are non-synonymous or chemically dissimilar across species. This binding pocket was consistently predicted by PrankWeb, MDockPep, and ClusPro and aligns with docking models and crystallographic data from *Bombyx mori* (PBAN/PBANR), *Homo sapiens* (NMU/NMUR), and Class A GPCRs more broadly [68,75,76,80]. Within the binding pocket, the conserved FXPRL peptide motif makes many contacts with conserved TM residues deep within the receptor. The entire peptide interacts with residues at the extracellular surface of the membrane, both in the TMs and ECLs, and a conserved β-turn in ECL2, the latter of which includes a conserved disulfide bond between cysteines in ECL2 and TM3 that is an essential structural feature for Class A GPCR activation [68,81,88,89]. Peptide N-termini also bind unique and often divergent amino acids in the ECLs, especially ECLs 2 and 3. Such a pattern of ligand-receptor interactions would allow the functional separation of peptides with unique binding properties and effects without compromising receptor recognition of the shared FXPRL peptide core motif.

PBAN’s extended 13 amino acid N-terminus interacts with several unique receptor residues, especially in the receptor’s N-terminal domain and distal ECL2 and 3. This would serve to differentiate the PBAN binding from that of the much shorter γ-SGNP and permit functional divergence of the two peptides. The important role of ECL3 in peptide recognition and receptor activation has been demonstrated in *Heliothis zea*, where replacing PBANR ECL3 with that of *D. melanogaster* PK1R (DHR) and vice versa disrupted ligand recognition [24]. γ-SGNP shares three receptor binding sites with PBAN and has just three unique binding sites, one each in TM3, TM4, and the N-terminal segment of ECL2. Hugin binds six unique receptor residues, four of which are non-synonymous with amino acids at the same location in PBANR. Hugin also shares six binding loci with PBAN: five of which are located in ECLs and four comprising non-synonymous amino acid substitutions. The divergence of binding sites between hugin and PK2_1 from their homologs γ-SGNP and PBANR supports coevolution of peptide and receptor in *D. melanogaster*. Binding of hugin to loci occupied by PBAN and the sparse colocalization of γ-SGNP and PBAN binding sites on PBANR suggests that loss of PBAN in *D. melanogaster* removed constraints on hugin binding, permitting interactions with residues in locations that differentiated between PBAN and γ-SGNP in *A. aegypti*.

Docking models highlight further complexity of receptor-ligand interactions. Individual peptide residues often contact multiple receptor sites, and vice versa. As described previously, peptide N-terminal residues engage unique receptor sites in the ECLs, many of which are non-synonymous between the two species, while the FXPRL core motif anchors the peptide to conserved sites in TMs and proximal ECLs. The variable “X” residue in the FXPRL motif (e.g., FAPRL in PBAN, FSPRL in γ-SGNP, FKPRL in hugin) has distinct receptor binding interactions in full-length and isolated core motif docking predictions and may further provide additional differentiation between peptides binding the same receptor as is the case in *A. aegypti*. Consistent with this prediction, the “X” residue of each peptide is conserved within phylogenetic groups. For example, FAPRL predominates in PBAN sequences of basal Diptera and many Coleoptera and Hemiptera (except Sternorrhyncha in the latter), whereas FSPRL is the most common PBAN core sequence in Lepidoptera (data not shown).

One feature of FXPRLamide neuropeptides that could not be included in the present study is the effect of the C-terminal amidation on peptide structure and receptor docking. Many studies have demonstrated that the amide group is critical to ligand recognition and activation, however, its binding interactions with the receptor have been investigated only in one species, the moth *Bombyx mori* [75]. The human FXPRLamide homolog NMU-25 amide group has also been shown to bind its cognate receptor and to be necessary for activation [76,80,90]. However, it has been difficult thus far to generate structural models of post-translational modifications using x-ray crystallography or *in silico* methods. However, new technologies are promising and future studies will avail of these methods to develop a more complete picture of FXPRLamide receptor binding [91–93].

In addition to sequence divergence, differences in peptide secondary and tertiary structure provide a basis for differentiating between similar peptides due to disparate binding and receptor activation properties. Structural models of PBAN consistently adopt rigid α-helical structures in the N-terminal and core motif regions. Docking predictions place the N-terminal along the inner surfaces ECL2 and ECL3 where unique binding sites are located, which could increase binding affinity and stability of PBAN [94,95]. The core FAPRL motif resides deep in the binding pocket near the base of ECL1, its helical structure likely created due to helix-promoting residues Ala and Pro [82,83]. In contrast, γ-SGNP lacks a long N-terminus, does not form α-helices and shows greater conformational variability and placement with the receptor in structural models and docking predictions. Binding predictions for hugin to PK2_1R are similar, suggesting that the compactness and reduced secondary structure of these short peptides might facilitate dynamic binding profiles. Influences on receptor specificity and activation might permit modulation of receptor signaling as demonstrated in other species [24,27,30,34,58,96]. This could occur via a mechanism such as ligand bias, in which ligands elicit distinct conformational changes in the receptor that that predispose interaction with different downstream pathways, conferring functional selectivity [97–101]. Functional studies confirm that the C-terminal β-turn-forming FXPRL core motif alone is sufficient for PBANR activation [70–72,102]. In this study, docking predictions for isolated core motifs did not align with the core motifs of full-length peptides. For example, in full-length PBAN, the Phe of the core FAPRL motif makes extensive contacts with ECL1; in contrast, the isolated core motif is shifted towards the center of the binding pocket, reducing ECL1 engagement. This suggests that although the isolated FXPRL can bind and activate its receptor experimentally, it may have additional binding and perhaps functional properties in the context of the full peptide. Forcing alignment of the core motif of a full PBAN peptide to an isolated FAPRL motif reveals a surprising feature of peptide tertiary structure that also impacts peptide-receptor interactions. A “proline kink” made up of two Pro residues in the PBAN peptide backbone located between the N terminus and core motif redirects the N-terminal helix orthogonally, passing through the ECLs of the receptor. A proline kink is frequently associated with α-helices, producing a bend that facilitates packing with other helices as in transmembrane proteins [103,104]; in other types of proteins, conserved Pro residues influence folding kinetics [105]. In the present study, when PBAN is appropriately docked, the core motif is oriented so that the kink directs the N-terminus through the center of the receptor without clashing with the ECLs. Forced alignment of the appropriate isolated core motifs with full γ-SGNP and hugin peptides, both containing a single proline in the same location as in PBAN, also reorients their N-termini into the ECLs rather than along ECL surfaces. This further supports the conclusion that while the FXPRL motif is a key component of peptide recognition and binding serves as a key interaction for receptor activation, it operates in a broader context provided by secondary and tertiary structure of the full peptide. These additional layers of interaction likely contribute to functional divergence of peptide variants within and across species.

## ACKNOWLEDGEMENTS

SMF thanks the WVU Department of Biology for supporting this work.

## Supporting Information

**S1 Table. Percent identity of PBANR and PK2_1R GPCRs by receptor domain.** N-terminal extracellular region-EC N term; transmembrane regions-TM 1-7; intracellular loops; ICL 1-3, extracellular loops; ECL 1-3, C-terminal intracellular region; IC C term.

**S2 Figure. Alignment and comparison of full receptor sequences and binding by FXPRL neuropeptides. *A****.aegypti* γ-SGNP and PBAN peptides binding to the PBAN receptor, *Drosophila* hugin binding to the PK2_1 receptor as determined using ClusPro. *Bombyx mori* PBAN binding to PBANR and *Homo sapiens* NMU/NMS peptides to NMUR1 and NMUR2 [75, 76, 80]. Ligand binding sites are shaded; underline indicate transmembrane regions of the receptors.

**S3 Figure. Receptor amino acids bound by the same residue in all three peptides and in both species.** Color = which peptide amino acid: grey- N terminus, magenta- F, blue- X (A, S or K), green- P, red- R, yellow- L.

**S4 Table**. **Combined contacts between individual peptide amino acids with regions of their cognate receptors.** N-terminal- all peptide residues before the C-terminal core motif FXPRL.

**S5 Figure. All valid ClusPro generated docking models using peptide models produced by PepFold4**. The receptor ECL1 is removed to provide a clearer view of the peptide. A-G, PBAN to *A. aegypti* PBANR. H-N, γ-SGNP to *A. aeg* PBANR. O-T, hugin to *D. mel* PK2_1R.

**S6 Figure**. **Comparison of docking predictions for γ-SGNP and hugin isolated core sequences (yellow) and full peptides (purple) before (γ-SGNP A, B; hugin E,F) and after forced alignment of the core motif (γ-SGNP C, D: hugin G, H).** Arrowheads in all figures indicate a proline residue introducing a kink in the peptide backbone. I, H. Consensus of docking predictions for the isolated FXPRL and full peptide core motifs with their cognate receptors. G- γ-SGNP with PBANR, H- hugin with PK2_1R. Colors indicate residues of each peptide: magenta, Phe (F); blue, A; green, Pro (P); red, Arg (R); yellow, Leu (L); grey- N terminal (full PBAN only). Only receptor ECL and TM binding regions are shown.

## Notes

### Competing Interest Statement

The authors have declared no competing interest.

